# Analyzing coordinated group behavior through role-sharing: A pilot study in female 3-on-3 basketball with practical application

**DOI:** 10.1101/2024.09.16.612561

**Authors:** Jun Ichikawa, Masatoshi Yamada, Keisuke Fujii

## Abstract

A group often shares a common goal and accomplishes a task that is difficult to complete alone by distributing roles. In such coordination, the nonverbal behavior among three or more members complicates the explanation of the mechanism due to complex and dynamic interactions. In cognitive science, a crucial role is indicated: to intervene moderately with others and adjust the whole balance without interrupting their main smooth interactions, using an experimental task. The findings suggest that resilient helping actions in the third role support to coordination. These actions are similar to off-ball movements in team sports, which involve an on-ball player and have recently been the focus of sports science because their characteristics are not represented in common statistical data, such as a shooting success rate. Hence, a new perspective for discussing coordination has emerged, as existing theories, such as synchronization—where movements between players are spontaneously matched and organized—cannot explain the mentioned role. However, there is a lack of investigation and discussion regarding whether these findings are applicable to real-world activities. Therefore, this study applied the experimental findings to the field of sports. We developed a 3-on-3 basketball game in which the offensive role of intervention decision and adjustment is key for winning and introduced it to the practice of a female university team as a pilot study. Participants repeatedly engaged in the mini-game, and the playing was compared before and after receiving tips of this role. Consequently, in the bins of the relatively large distance between the participant required to the relevant role and each defensive player, the frequencies after receiving these tips were significantly higher. Furthermore, the winning rate on the offensive team improved temporarily; however, the effects were not maintained. These suggest that spacing skill, which maintains reasonable distances from the other players, creates favorable situations for coordination. This study may bridge the gap between controlled experiments and real-world applications and make an educational contribution; it may recommend practice design for the acquisition of spacing skill related to the crucial role.

## 1 Introduction

A group often shares a common goal and accomplishes a task that is difficult to complete alone by distributing roles. This is explained by common opinions or suggested through experiments and simulations in many previous studies (e.g., (1-4)). That is encapsulated by the saying “Two heads are better than one” and defined as planned coordination (coordination) in psychology and cognitive science (5). For coordination, members share some roles considering the workloads and task specifications. Distributed cognition theory explains that an overall group function works through interactions among subsystems and between subsystems and environments, in which each subsystem plays a role (6,7). The internal and external resources are properly distributed, and each member’s workload decreases. Furthermore, cognitive interactions among roles based on different perspectives lead to problem-solving and discovery of various strategies (e.g., (8-12)). Team sports, in particular, are typical examples of the benefits of role-sharing by multiple on- and off-ball players (e.g., (13-16)). This is required to achieve a common goal and high performance under rules, time constraints, and defensive pressures. Team sports often involve three or more members and aim to achieve a common goal through complex and dynamic interactions. Complexity indicates that explaining the coordination mechanism is more complicated than for a pair because relationships among members diversify, such as role-sharing, and are not interpreted by one-dimensional behaviors such as leader-follower and approach-avoidance (17,18). Dynamics suggests nonverbal and time-series features that develop strongly over a short period, such as body movements (19,20). Cognitive science focuses mainly on higher-order information processing, and coordinated behaviors among three or more members are not fully investigated. Considering that physical interactions are primitive, and that a group of three or more members is often observed in real-world activities, it is important for cognitive science, which discusses social intelligence, to investigate complex and dynamic coordination (17).

This study focuses on a crucial role in coordinated behavior of a triad through role-sharing, which is indicated using an experimental task in a cognitive science study (21). We aim to bridge a gap between experimental findings and real-world applications and contribute to the understanding and developing of coordination mechanisms.

As related works, a sports science study recorded 5-on-5 basketball games and identified the coordinated defensive structures of role-sharing according to emergent situations by a top-level male university team in Japan (1). Recently, machine learning studies have predicted shooting success rate and evaluated coordination leading to goals by classifying and extracting features of physical interaction structures with role-sharing using datasets in basketball and soccer (e.g., (22-24)). Furthermore, network science studies have identified indices of centrality in network models where nodes and links are regarded as players and passes in soccer, representing team-specific strategies and key players related to robust coordination (e.g., (25-27)). Meanwhile, collective behavior studies observed coordinated hunting and found that the role-sharing of chasing and blocking naturally emerged (e.g., (28,29)). Additionally, multi-agent simulation using deep reinforcement learning can replicate this process. In the simulation, agents perceive, observe the environment, including the prey, and learn the optimal behaviors to maximize their rewards (30). The findings in sports science, machine learning, network science, and collective behavior suggest that role-sharing creates favorable situations for coordination. In principle, these studies aim to understand the physical interaction structures in coordination. However, the information processing underlying these characteristics has not been fully discussed, and educational applications have not been considered.

Notably, although not competitive, a previous cognitive science study investigated a crucial role in coordinated behavior of a triad through role-sharing using an experimental task (21). This indicates that the role of intervening moderately with other roles and adjusting the whole balance was related to high task performance. Such resilient helping actions are an important factor for successful defensive coordination in team sports and effective team building in business and military organizations, not only in experimental tasks (e.g., (1,31-34)). This role is also required not to interrupt their main smooth interactions. The third role needed to decide whether to intervene according to the situations. A new perspective for discussing coordination has emerged because existing theories in cognitive science and sports science, such as synchronization (e.g., (35-38))— where movements between players are spontaneously matched and organized—cannot explain this role. The findings suggest that adjustment to create favorable situations without interrupting others supports coordination. This concept is similar to off-ball movements in team sports involving an on-ball player. For example, in basketball, when an on-ball player is surrounded by defensive opponents, another offensive player approaches and directly receives a pass for helping. Off-ball movements in basketball and soccer have recently been the focus of sports science, as shown by distances between players and those with the goal. These contain valuable information on coordination and are not represented in common statistical data, such as shooting success rate (e.g., (13,16,39,40)). The previous study (21) confirmed the crucial role in coordination; however, there is a lack of investigation and discussion regarding whether these findings are applicable to real-world activities.

Therefore, we applied the experimental findings to the field of sports as a pilot study. This study focused on team sports that must achieve a common goal and high performance within some constraints. We used 3-on-3 basketball in which the coordination by off-ball players is essential for winning, as mentioned above. Furthermore, it is easy to record group behavior because of the relatively small number of players (three on each team) and the court size. Additionally, it has recently attracted worldwide attention, as evidenced by its inclusion as an official Olympic event. Meanwhile, few studies have used 3-on-3 basketball to discuss coordination in terms of cognitive science. The purpose of our study was to investigate the influence of the role of intervention decision and adjustment on coordination in 3-on-3 basketball. We developed a mini-game in which the relevant offensive role is key and introduced it to the practice of a female university team. The players repeatedly engaged in the game. After the first half, the offensive team received tips on coordination focusing on this role. The team performance and playing related to the relevant role were quantitatively compared before and after receiving these tips. According to the purpose, this study has set the following hypothesis: after receiving the tips, the team performance improves, and the appropriate role executions are observed.

Our study connects cognitive science and sports science because this pilot study includes investigation of information processing underlying complex and dynamic coordination, which has not yet been fully discussed. The quantitative analysis may recommend practice design on playing that helps an on-ball player and creates favorable situations in the fields of sports. Bridging the gap between controlled experiments and real-world applications is a challenging endeavor. These findings may also offer implications for further developing 3-on-3 basketball itself. Next, we explain the details on this practice.

## 2 Methods

### 2.1 Participants

Six female students on the university basketball team, of which the second author is the head coach, participated in this practice. They usually play 5-on-5 in the official game. This team was affiliated with the third division of the Tokai area league in Japan, practiced regularly for about two hours each time three or four times a week, and played approximately 30 games per year including practice matches. The participants included five regular players and sixth man. Sixth man means a first substitute player. Their averages of age, height, and basketball experience were 19.17 age (*SD* = 0.90), 165.08 cm (*SD* = 5.88), and 10.58 years (*SD* = 2.70), respectively.

The participants were divided into the offensive and defensive teams. The head coach conducted the team compositions to make their abilities competitive based on their profile data of age, dominant hand, height, basketball experience, and current and previous positions (see the details on the offensive and defensive teams in Supplementary Materials). The second author has 25 years of coaching experience and holds a certified B-level coaching license from the Japan Basketball Association. The record includes leading a university team to promotion to the first division of the Tokai area league and participation in national tournaments in Japan. The participants on each team were fixed. This study statistically compared the offensive team performance and playing related to the role of intervention decision and adjustment before and after receiving the tips of this role. Hence, we planned this practice based on the policy of eliminating factors other than these tips as much as possible. Meanwhile, it should be kept in mind that we applied the experimental findings to the field of sports as a pilot study.

### 2.2 Informed Consent

We explained how we would video-record and collect data. When explaining, not all the participants were informed of the details on the procedures in this practice. They were not made aware of the playing related to the role of intervention decision and adjustment, receiving the tips of this role to the offensive team, and comparing the offensive team performance and playing before and after receiving these tips. The procedures were also based on the policy of eliminating factors other than the tips. Written informed consent was obtained from all the participants. This study was approved by the ethics and safety committee of Shizuoka University and Tokoha University, to which the first author and the participants were affiliated. Our study was conducted following these regulations. According to the informed consent, not all images in the manuscript contain individual identifiers.

### 2.3 Developed 3-on-3 basketball game

Figure 1 shows the diagram of the developed mini-game in which the circles and squares represent the offensive and defensive players, respectively. Players from #1 to #3 on each team are the same participants in all the games (trials). Offensive #3 is required to play the role of intervention decision and adjustment focused on in this study. The winning condition for the offensive team is that within a 15-second time limit, someone draws a defensive player and a shooter is unmarked in open space. Conventionally, the defensive team must play without satisfying it. To investigate coordination, the above condition is defined independently of individual skills. The team performance reported in the Results section is consistent with this definition.

**Figure 1.**
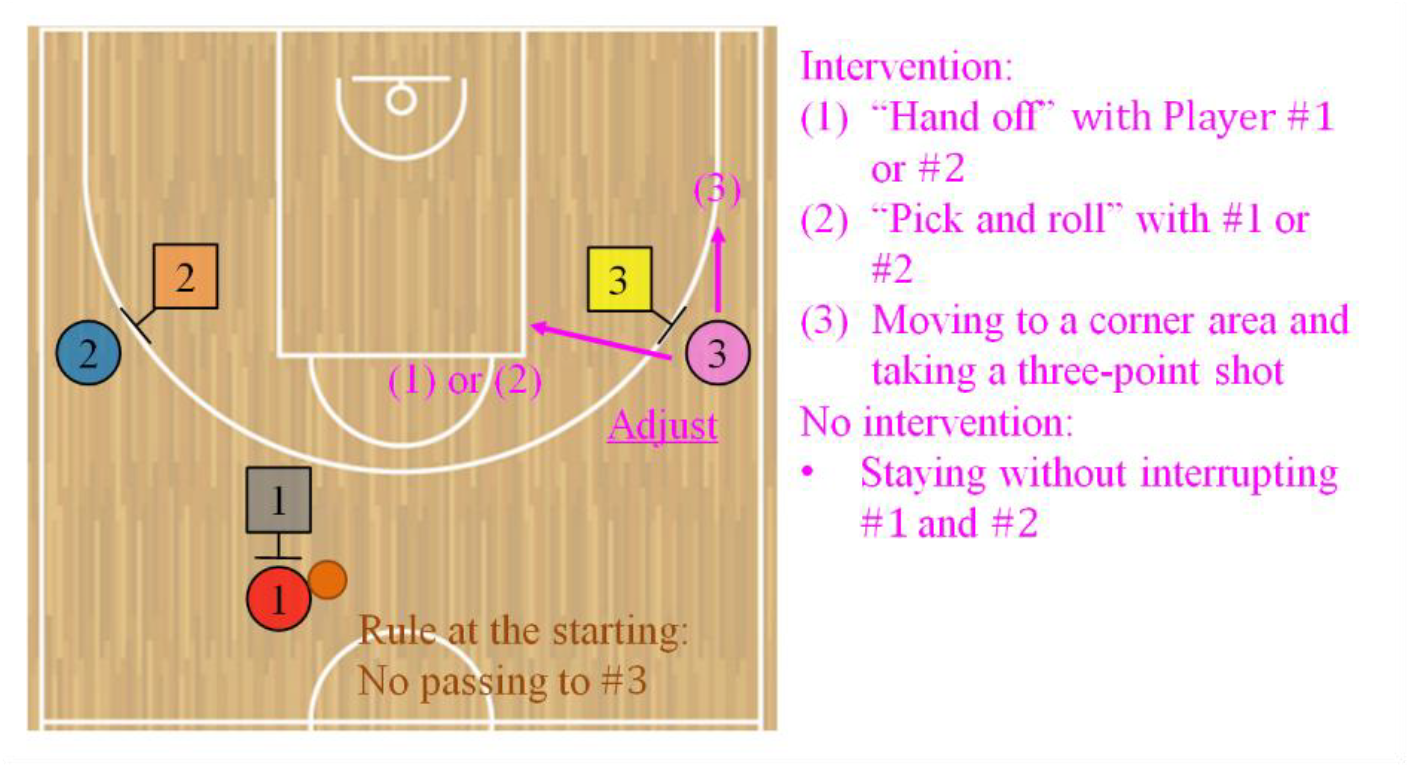
Diagram of the developed 3-on-3 basketball game in this study. In this mini-game, the circles and squares represent the offensive and defensive players, respectively. The winning condition for the offensive team is that within a 15-second time limit, someone draws a defensive player and a shooter is unmarked in open space. At the start of the game, the ball handler of #1 is fixed, and the player begins without directly passing to #3. The starting position of each offensive player is assigned (see the Developed 3-on-3 basketball game section to confirm the other rules). For the offensive team to win, #3 is a key player; it is required to intervene with the other players, adjust the 2-man game, and create favorable situations, such as (1) handing the ball off directly with #1 or #2 (“hand off”), (2) working as a wall to interrupt the defensive player (“pick and roll”), and (3) moving to a corner area and taking a three-point shot. Conversely, it is crucial not to interrupt #1 and #2 without forcibly intervening according to the situations.

In the mini-game, we set some rules to guide the same situation at the start of the game and compare the playing before and after giving the tips about the role of intervention decision and adjustment. The ball handler #1 is fixed and the player begins the game without directly passing to #3. The starting position of each offensive player is assigned to maintain a certain distance to observe the coordination process in which this role involves the other players. On the defensive team, #1 and #2 confront offensive #1 and #2 and their starting positions are voluntary. The start position of defensive #3 is also voluntary. Just before the game, the 2-man game are randomized on the right (Figure 1) or left side of the basket goal by instructions to eliminate the influence of the dominant hand. The starting positions on the offensive team are called the two-guard and also used in a 5-on-5 game. For the offensive team to win, it is important that #3 intervenes the other players, adjusts the 2-man game, and creates favorable situations, such as (1) handing the ball off directly with #1 or #2 (“hand off”), (2) working as a wall to interrupt the defensive player (“pick and roll”), and (3) moving to a corner area and taking a three-point shot. Conversely, it is also key not to interrupt #1 and #2 without forcibly intervening according to the situations. The offensive coordination required in this game is fundamental. However, the head coach mentioned that the participants could not implement such interaction in official games at that time.

### 2.4 Environment and Procedures

The upper part of Figure 2 shows the environment of the mini-game conducted in the university gymnasium. The court area, including the vertical endline and the area under the basket goal (“paint area”), followed the official size of 3-on-3 and 5-on-5 (41,42), except for the horizontal sideline. A digital countdown timer was used to watch the remaining time (mollten Corp., UX0110). The participants regularly practiced on this court size in the gymnasium. Hence, factors such as the court area, lighting, and flooring were unlikely to negatively influence the offensive team performance and group behavior. The mini-games were recorded from a bird’s-eye view on the stage using only one video camera, as shown in the lower part of Figure 2 (Sony Corp., HDR-CX680). As they played all the trials within the camera’s field of view, there were no issues with tracking their positions by image processing technology as below. We preliminarily prepared another video camera; however, no equipment problems occurred during this practice. Consequently, these recordings were excluded for this analysis.

**Figure 2.**
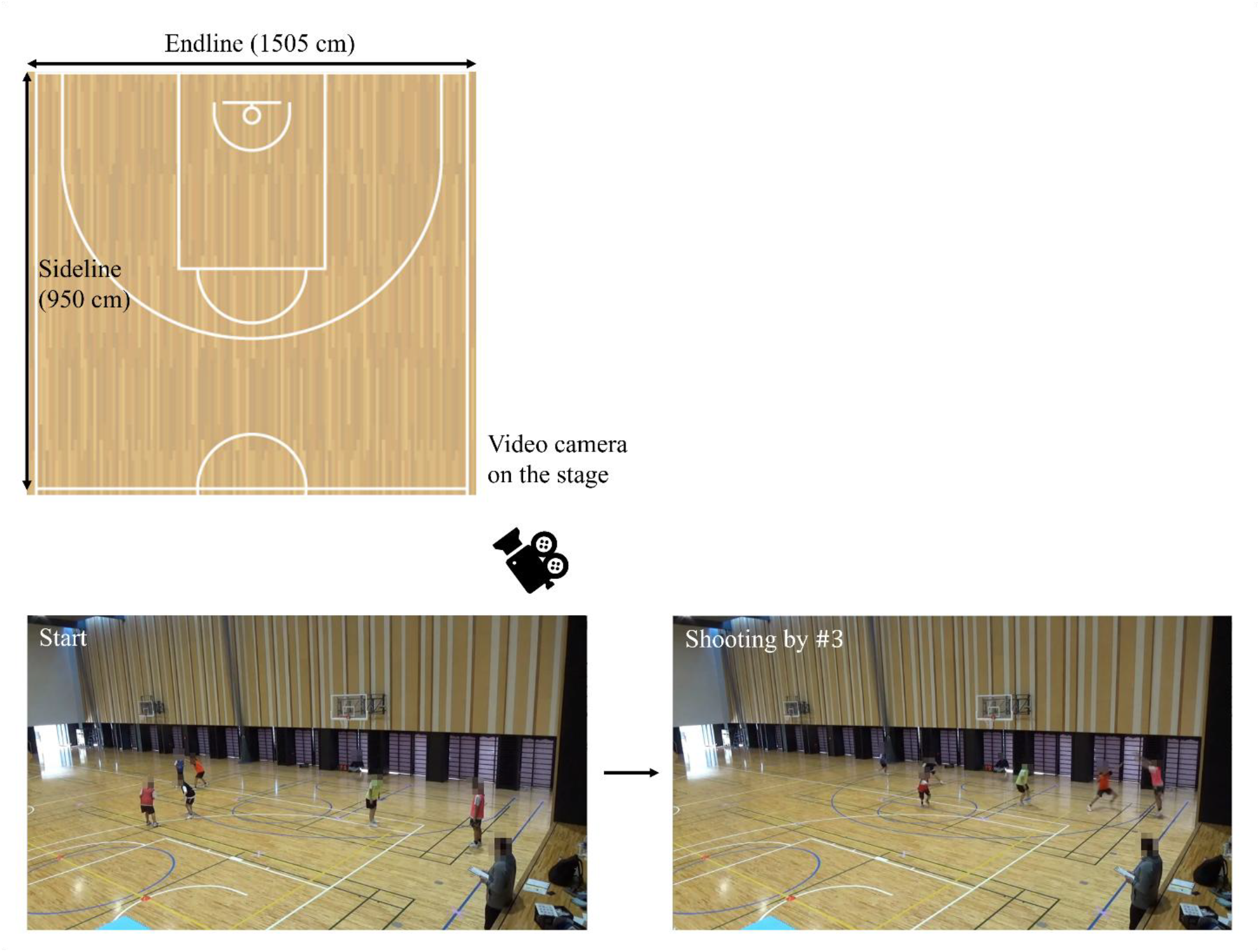
Environment in this practice. We conducted the developed 3-on-3 basketball game in the university gymnasium. The court area, including the vertical endline and the area under the basket goal (“paint area”), follows the official size in 3-on-3 and 5-on-5 (41,42), except for the horizontal sideline. All the images in the mini-games were recorded from a bird’s-eye view on the stage using only one video camera by the first author. After receiving the tips about the role of intervention decision and adjustment, #3 required to this role, wearing a pink bib as shown in Figure 1, moves to a corner area while maintaining a reasonable distance from each defensive player and aims to take a three-point shot, as shown in the lower images. We explained how we would video-record and collect data. Written informed consent was obtained from all the participants. According to the informed consent, all the images are shown while blurring some parts to avoid identifying individuals.

Regarding the procedures, the experimenter briefly announced the practice schedule (see these details in Supplementary Materials). Subsequently, the participants warmed up and received colored bibs for individual identification (Figure 1). The experimenter then explained the team assignments and mini-game rules. At this point, the first author did not instruct the offensive team on the role of intervention decision and adjustment and playing. The three practice trials, including a trial by the timer error, were conducted and they could confirm the rules. Subsequently, in the first half as the pre-test, three sessions comprising seven trials per session were conducted for a total of 21 trials. The interval between the trials was approximately 30 seconds, with about a minute between the sessions. The university team regularly played a 5-on-5 game with each quarter for 10 minutes with the two minutes intervals for a total of 40 minutes across the four quarters. In comparison, the total of playtime in this practice was a maximum of approximately 10 minutes. Hence, fatigue was unlikely to negatively influence the offensive team performance and group behavior. Furthermore, we carefully checked the participants just before restarting the mini-game to ensure no fatigue. After the first half, a guest coach who understood the purpose of this study gave each team tips based on the observations of the games. The coach was a professional player in Japan and is currently a head coach of a junior youth team. In this practice, giving the tips that reflected the essence of our study to the offensive team was indispensable. Thus, we asked the guest who is superior to coaching to do so. The tips for the offensive team focused on coordination related to the role of intervention decision and adjustment. Meanwhile, the defensive team received the general tips on help defense, excluding participant- and game-specific strategies. Table 1 represents details of the tips given to each team. While one team received them, another waited at a distance to prevent information gathering. Generally, it is common for a team to keep strategies hidden from the opponent. The purpose of this study was to investigate the influence of the tips on the offensive team performance and playing related to this role. Hence, we planned this practice based on the policy of eliminating factors other than giving the offensive tips; the mentioned procedure was appropriate. Subsequently, the second half as the post-test was conducted similar to the first one.

**Table 1.** Tips to the offensive and defensive teams by the guest coach.

| Team | Tips |
| --- | --- |
| Offense | • #3 checks the position of defensive #3. <u>When defensive #3 helps, #3 moves to where the driving offensive player (#1 or #2) can see #3, such as a corner area (“deep corner”), so that #3 can easily take a three-point shot in open space.</u><br>• The very first thing #1 and #2 should do is to play a 2-man game where #1 or #2 drives, that is, dribbles toward the area under the basket goal (“paint area”). If #1 pretends to cooperate with #2 working as a wall to interrupt the defensive player (“pick and roll”) and drives to the other side of the wall (“reject”), defensive #3 has to move to help. <u>Then, #3 moves to the corner along the three-point line taking advantage of a wide space.</u><br>• However, as an exception, in a case where #3 is in a high position (2-man game position), if defensive #3’s back is visible from #3 and defensive #3 goes for help, <u>#3 should dive to the paint area because moving along the three-point line and receiving the pass take time.</u><br>• <u>#3 should not move too much. #3 should carefully check the defensive players’ positions while staying around:</u> (1) the wing position (45-degree position on the three-point line with the goal as the origin), (2) the position near the three-point line changing from straight to curved, or (3) the corner position. |
| Defense | • If #3 does not help, the space near the goal becomes open for a shot, so #3 has to move to help. If offensive #3 receives the ball, #3 should mark offensive #3 immediately. The defensive players need to check where the ball is and it makes easier to help. If not, the offensive player can take an uncontested shot with a running jump (“lay-up shot”).<br>• Force the offensive player to make a long, arching pass in order to buy time for getting the defensive ones ready. The defensive players’ keeping their hands up will force the offensive one to make an arching pass, creating time for them to close the distance with the offensive one (“closeout”). |

## 3 Results

### 3.1 Team Performance

This study counted the winning numbers of the mini-games for the offensive team to compare them between the first and second halves. Those indicated in the first half for a total of 21 trials from Sessions 1 to 3 and in the second one from Sessions 4 to 6 were 9 and 11, respectively. Table 2 represents a cross-tabulation table of the event and performance. A chi-square test indicated no significant relationship between them, and the effect size φ was small (*χ*^2^(1) = 0.095, *p* = 0.757, *φ* = 0.048). Meanwhile, Figure 3 shows the progression of the winning rates on the offensive team over the sessions. The rates of Sessions 1 and 2 in the first half were at the chance level, indicating four wins in the seven trials; thereafter, it decreased in Session 3. Notably, in the second half, it recovered in Session 4 and drastically improved in Session 5, indicating six wins; however, it significantly decreased in Session 6. It indicated that after receiving the tips about the role of intervention decision and adjustment, the offensive team performance improved temporarily but was not maintained.

**Table 2.** Cross-tabulation table of the event and offensive team performance.

| Event | Offensive team performance |  |
| --- | --- | --- |
|  | Win | Loss |
| First | 9 | 12 |
| Second | 11 | 10 |

**Figure 3.**
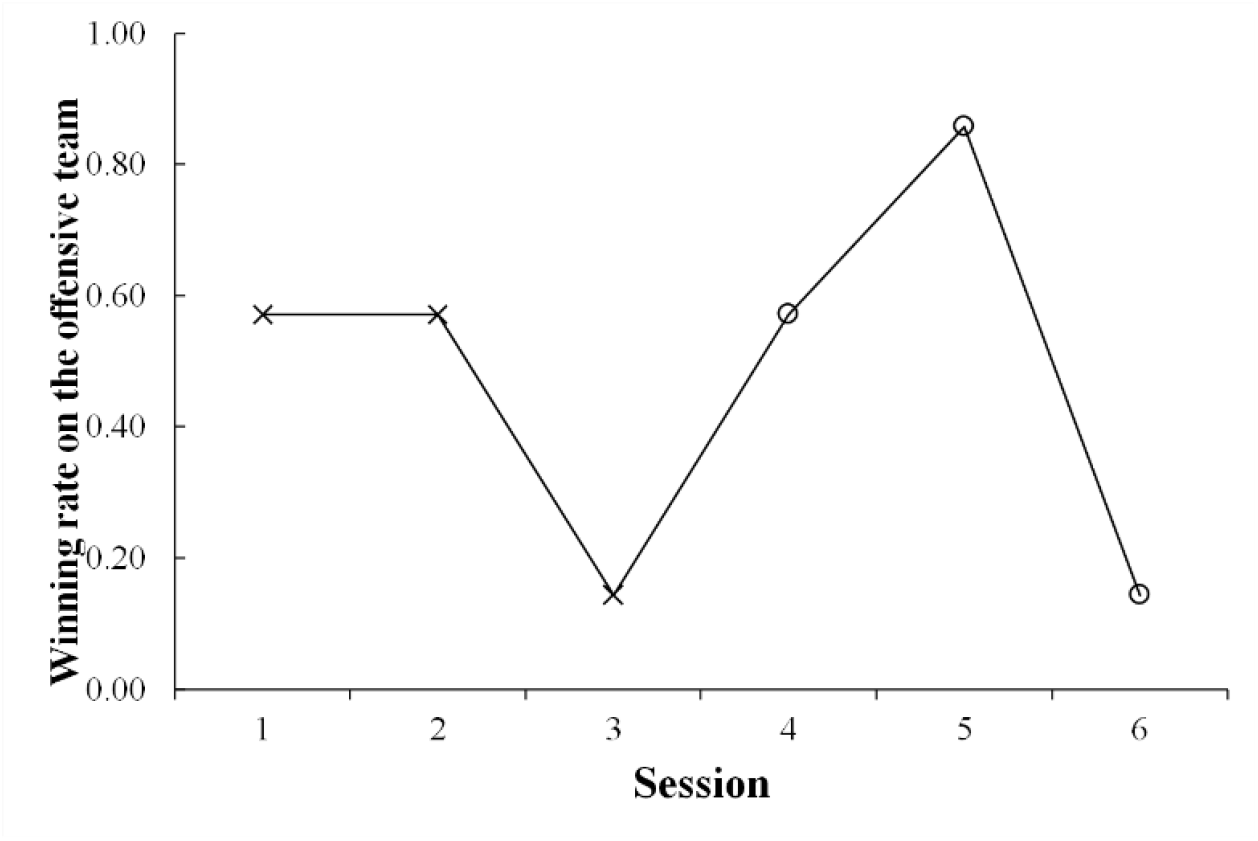
Progression of the winning rates on the offensive team over the sessions. Each session consists of the seven trials. After Session 4, given the tips about the role of intervention decision and adjustment, the team performance improves temporarily but is not maintained.

In summary, giving the tips about the role of intervention decision and adjustment did not fully influence the offensive team performance. The main reason for decreasing the winning rate in the final session was that the defensive team had established an effective countermeasure, as reported by the participants on the defensive team (see these details in the Discussion section). The offensive team might not achieve developed coordination based on the tips of this role because of some factors such as only a single practice and individual skills.

### 3.2 Analysis Procedures of Playing Related to the Offensive Role of Intervention Decision and Adjustment

We obtained time-series position data of each participant in the two dimensions, x- and y-components, through the image processing of a bird’s-eye view recording (Figure 2). This study compared the movements of offensive #3, required to the role of intervention decision and adjustment, as shown in pink in Figure 1, between the first and second halves using the dataset. The participants on the movies (20 fps, 1280 px × 720 px) were tracked using ByteTrack (43) based on YOLOX. The experimenter manually corrected false detections and undetected positives using the labeling platform Labelbox (44), and prepared the dataset from real coordinate transformations. The average of the absolute measurement errors was 3.165 cm × 2.102 cm, similar to those (2.3 cm × 4.6 cm) reported in the previous study (18) that investigated passing coordination by a triad in soccer. Considering the bin interval of 200 cm in the histogram used in the analysis explained below, the errors in our study were unlikely to negatively influence the results. Meanwhile, in this practice, it is difficult to detect a ball using image processing technology due to the recording environment and technical problems; this is future work.

This study analyzed (1) the distance (cm) between the offensive participant required to the role of intervention decision and adjustment and each defensive player, as shown in black, orange, or yellow in Figure 1, and (2) the distance between the offensive key player and each other participant, as shown in red or blue in Figure 1. We evaluated spacing skill to play this role. Spacing skill, which maintains reasonable distances with other players, is generally crucial for both offensive and defensive coordination in basketball (e.g., (13,14,16,45,46)). In the mini-game, if reasonable distances from the defensive players are maintained, the defensive pressure is reduced, which makes it easy to intervene moderately with the other offensive players according to the situations, as explained in Table 1. If those among the offensive players are maintained, it is easy to pass the ball to the relevant role, and the defense will break down. The characteristic is also represented by staying in place without interrupting the other offensive players, as shown in Table 1. This analysis calculated both indices of (1) and (2) for each time frame and made these histograms. Histogram analysis is a fundamental method to understand overall trends for investigating complex and dynamic coordination. It has been used in previous studies on group behavior (e.g., 4,47,48) because of its advantage in intuitively capturing the characteristics of continuous values. Histograms were generated in each trial, and the normalized frequencies were averaged for the first and second halves. A *t*-test was conducted to compare them between the first and second halves in each bin. It is expected that significant differences in the frequencies for the indices of (1) and (2) are confirmed because of giving the tips of this role. This study conducted the exploratory comparisons of reasonable distances between the participants because of analyzing complex and dynamic group behavior in the field of sports.

The previous studies (4,48) conducted the *t*-test in each bin of a histogram to statistically compare frequencies between different conditions. However, we should keep in mind the type I error by the multiple comparisons. Therefore, this study calculated the effect sizes (Cohen’s *d*) and powers (1-*β*) in all the bins for both indices of (1) and (2) to improve the validity of the statistical tests and carefully interpret these results. For calculating the powers, we set the significant level at 5% (α= 0.05), effect sizes, and sample size *n* (21 trials) in each half. We comprehensively evaluated the differences between the first and second halves in terms of the *p*-value, effect size, and power indices. The Bonferroni method was not applied to avoid because the numbers of the bins in both indices of (1) and (2) were large and it was easy to cause the type II error. Our policy was to avoid this problem by calculating the three indices and investigating the significant differences between the first and second halves from the different perspectives. The distances between the participants were analyzed using MATLAB R2021b. The statistical tests and the investigation of these validities were conducted using R 4.3.3 and G*Power 3.1, respectively. However, it is crucial to compare the playing related to the role of intervention decision and adjustment using applied statistical methods before and after giving the tips of this role; this is also future work.

### 3.3 Results of the Distance with Each Defensive Player

Figure 4 shows a time-series example of the distance (cm) between the offensive participant required to the role of intervention decision and adjustment and each defensive player in a trial. Figure 4 also shows the histogram in each session. The upper and lower parts represent those in the first and second halves, respectively. The horizontal and vertical axes indicate the bin and average of the normalized frequencies across the seven trials of each session, respectively.

**Figure 4.**
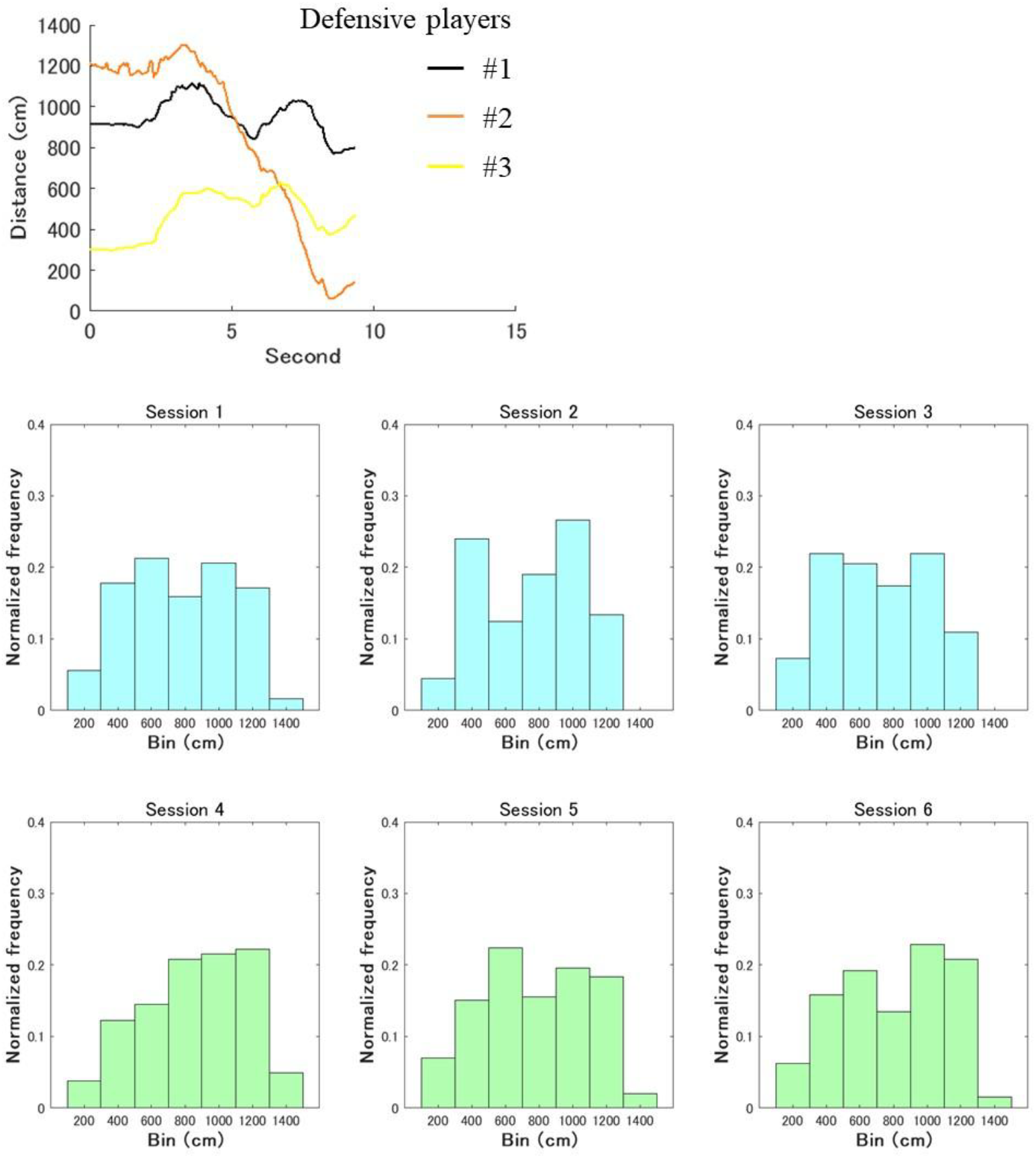
Overall characteristics of the distances (cm) between the offensive player required to the role of intervention decision and adjustment and each defensive participant. Its time-series example in a trial fluctuates up and down. Meanwhile, the upper and lower parts represent the histograms in the first and second halves, respectively. The horizontal and vertical axes indicate the bin and average of the normalized frequencies across the seven trials of each session, respectively.

In each bin, the average of normalized frequencies across the 21 trials in the first half from Sessions 1 to 3 was compared to that in the second one from Sessions 4 to 6. Notably, the *t*-tests confirmed the significant differences in the bins of 400 cm, 1200 cm, and 1400 cm, and these effect sizes ranged from medium to large while the powers were high (400 *cm*: *t*(20) = 2.739, *p* = 0.013, *d* = 0.894, 1 − *β* = 0.973; 1200 *cm*: *t*(20) = −2.269, *p* = 0.034 , *d* = 0.732, 1 − *β* = 0.891; 1400 *cm*: *t*(20) = −2.581, *p* = 0.018, *d* = 0.668, 1 − *β* = 0.829, Figure 5A–C). In the bin of relatively small 400 cm, the frequency in the second half was significantly lower than that in the first one. Conversely, in the bins of relatively large 1200 cm and 1400 cm, the frequencies in the second half were significantly higher than those in the first one. In the other bins, no significant differences were confirmed (see these details in Supplementary Materials).

**Figure 5.**
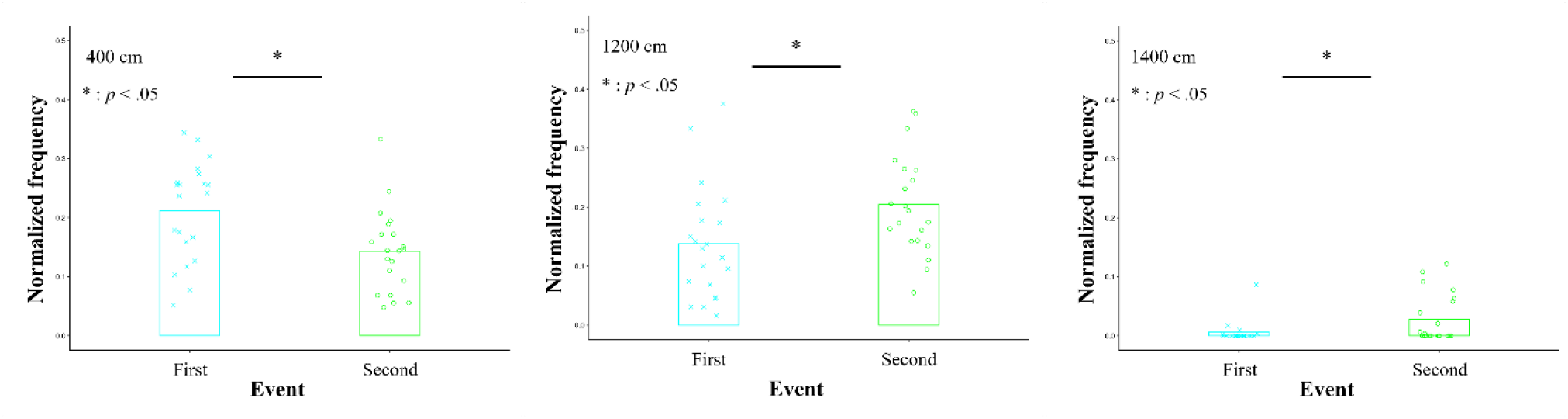
Averages of the normalized frequencies in the bins confirmed the significant differences between the first and second halves in the histograms of Figure 4. In the bin of relatively small 400 cm (A), the normalized frequency in the second half is significantly lower than that in the first one. Conversely, in the bins of relatively large 1200 cm (B) and 1400 cm (C), the frequencies in the second half are significantly higher than those in the first one.

### 3.3 Results of the Distance with Each Other Offensive Player

Figure 6 shows a time-series example of the distance (cm) between the offensive player required to the role of intervention decision and adjustment and each other participant in a trial. Figure 6 also shows the histogram in each session. The upper and lower parts represent those in the first and second halves, respectively. The horizontal and vertical axes indicate the bin and average of the normalized frequencies across the seven trials of each session, respectively.

**Figure 6.**
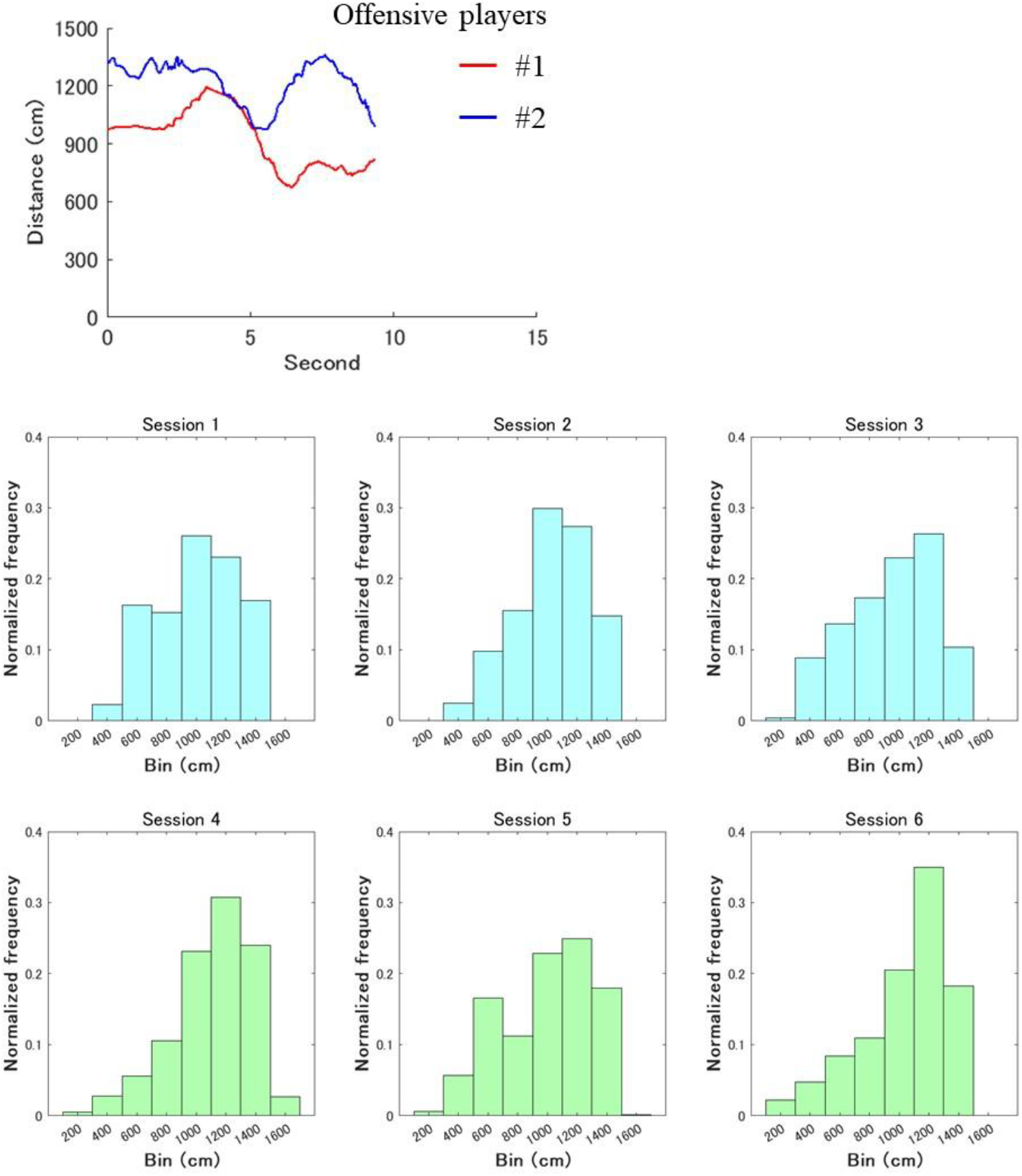
Overall characteristics of the distances (cm) between the offensive player required to the role of intervention decision and adjustment and each other participant. Its time-series example in a trial fluctuates up and down. Meanwhile, the upper and lower parts represent the histograms in the first and second halves, respectively. The horizontal and vertical axes indicate the bin and average of the normalized frequencies across the seven trials of each session, respectively.

In each bin, the average of normalized frequencies across the 21 trials in the first half from Sessions 1 to 3 was compared to that in the second one from Sessions 4 to 6. Considering the results of the *t*-tests, effect sizes and powers, the clear differences were not confirmed between the first and second halves. Although the effect sizes were medium and the powers were lower than those reported in the previous section, the *t*-tests indicated the significant trends of differences in the bins of 200 cm, 800 cm, and 1600 cm (200 *cm*: *t*(20) = −2.002, *p* = 0.059, *d* = 0.517, 1 − *β* = 0.616; 800 *cm*: *t*(20) = 1.856 , *p* = 0.078, *d* = 0.628, 1 − *β* = 0.782; 1600 *cm*: *t*(20) = −1.745, *p* = 0.096, *d* = 0.572, 1 − *β* = 0.703, Figure 7A–C). These showed a U-shape trend; in the bins of relatively small 200 cm and large 1600 cm, the frequencies in the second half tended to be significantly higher than those in the first one. Conversely, in the middle bin of 800 cm, the frequency in the second half tended to be significantly lower than the first one. In the other bins, no significant differences were confirmed (see these details in Supplementary Materials).

**Figure 7.**
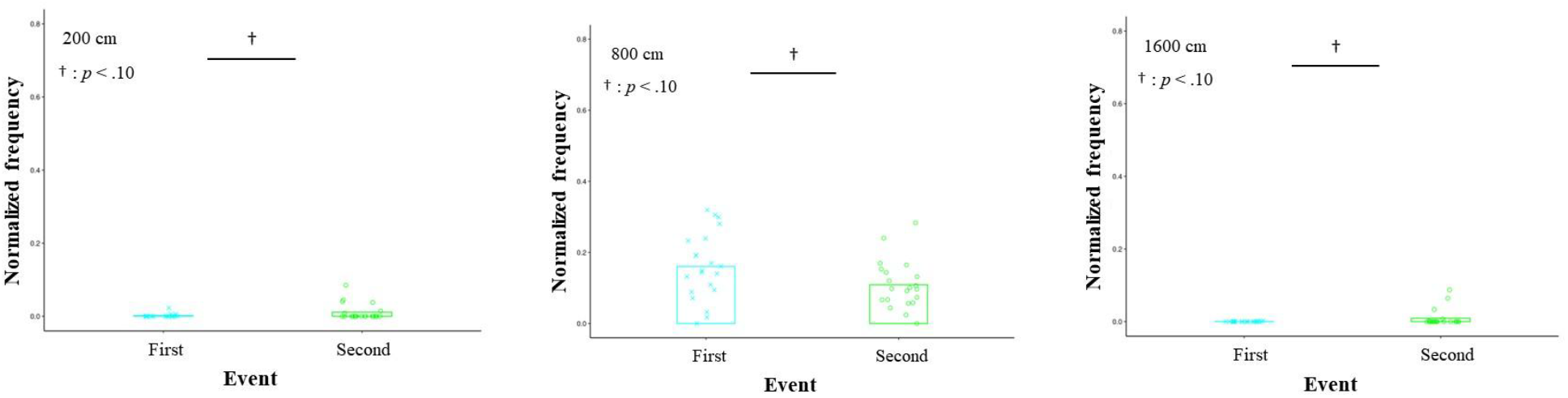
Averages of the normalized frequencies in the bins confirmed the significant trends of differences between the first and second halves in the histograms of Figure 6. In the bins of relatively small 200 cm (A) and large 1600 cm (C), the normalized frequencies in the second half tend to be significantly higher than those in the first one. Conversely, the frequency in the middle bin of 800 cm (B) in the second half tends to be significantly lower than that in the first one.

## 4 Discussion

This study developed the mini-game of 3-on-3 basketball, in which the offensive role of intervention decision and adjustment is crucial for winning. We introduced it to the practice of the female university team as a pilot study. The purpose of our study was to investigate the influence of this role on coordination. In this practice, the results confirmed that in the bins of the relatively large distance between the participant required to the relevant role and each defensive player, the frequencies after receiving the tips of the crucial role were significantly higher than before (Figure 5B,C). Conversely, in the bin of the relatively small distance, the frequency after receiving these tips was significantly lower than before (Figure 5A). Furthermore, the winning rate on the offensive team improved temporarily (Figure 3); however, the effects of receiving the tips were not maintained. Therefore, these results partially supported the hypothesis regarding the effects of giving the tips.

After receiving the tips, the participant required to the offensive role of intervention decision and adjustment maintained a reasonable distance from each defensive participant. This suggests that the spacing skill might create favorable situations for coordination, such as moving to the corner area and aiming to take a three-point shot, as explained in Table 1 (Figure 8). Although the effect sizes and powers were lower for the distance with the other offensive players compared to those with the defensive players, the notable trends were observed. In the bins of the relatively small and large distances, the frequencies after receiving the tips of this role tended to be significantly higher than before (Figure 7A,C). In the bin of the middle distance, the converse result was obtained (Figure 7B). Such a U-shape trend might indicate that the participant required to the relevant role also stayed in place without interrupting the other offensive participants, as explained in Table 1. However, these discussions should be investigated in more detail. The previous studies in basketball suggested that offensive interactions, involving ball movement to an opposite side and maintaining a reasonable distance with a defensive player, create a gap with the opponent (16,45). These lead offensive shooting opportunities in open space. In this practice, running to the corner area can serve as a starting point for moving the ball to the opposite side. This enabled the participant required to the crucial role to take a three-point shot in open space. Therefore, such coordination increases the countermeasures for the defensive team, making it difficult to anticipate the next playing. Spacing skill, including a reasonable distance with the goal, is related to high success rate of offensive shooting and defense in basketball. Expanding space within one’s own team and limiting it for the opponent team develop successful opportunities. These variables are key to predict team performance using statistical and machine learning models (e.g., 13,14,16,45,46). It is also essential to create favorable situations in soccer other than basketball and influences strategies (e.g., 49,50). The previous study (49) confirmed that the experimenter manipulated space to play in a soccer game and its limitations influence strategic behaviors. Another study (50) quantified and visualized the occupied space of each soccer player using applied geometric analysis with Voronoi diagrams. These results suggest that dynamical adjustment of defensive space according to situations limits opponent passing options. Although the types of team sports and methods used in these studies differed from those used in this study, the findings are similar with ours. Meanwhile, many previous studies mention above analyzed professional teams. Hence, our results from the female university team, which was affiliated with the third division league in Japan and did not have a high skill level, are valuable. Therefore, this study provides new insights into spacing skill related to the role of intervention decision and adjustment.

**Figure 8.**
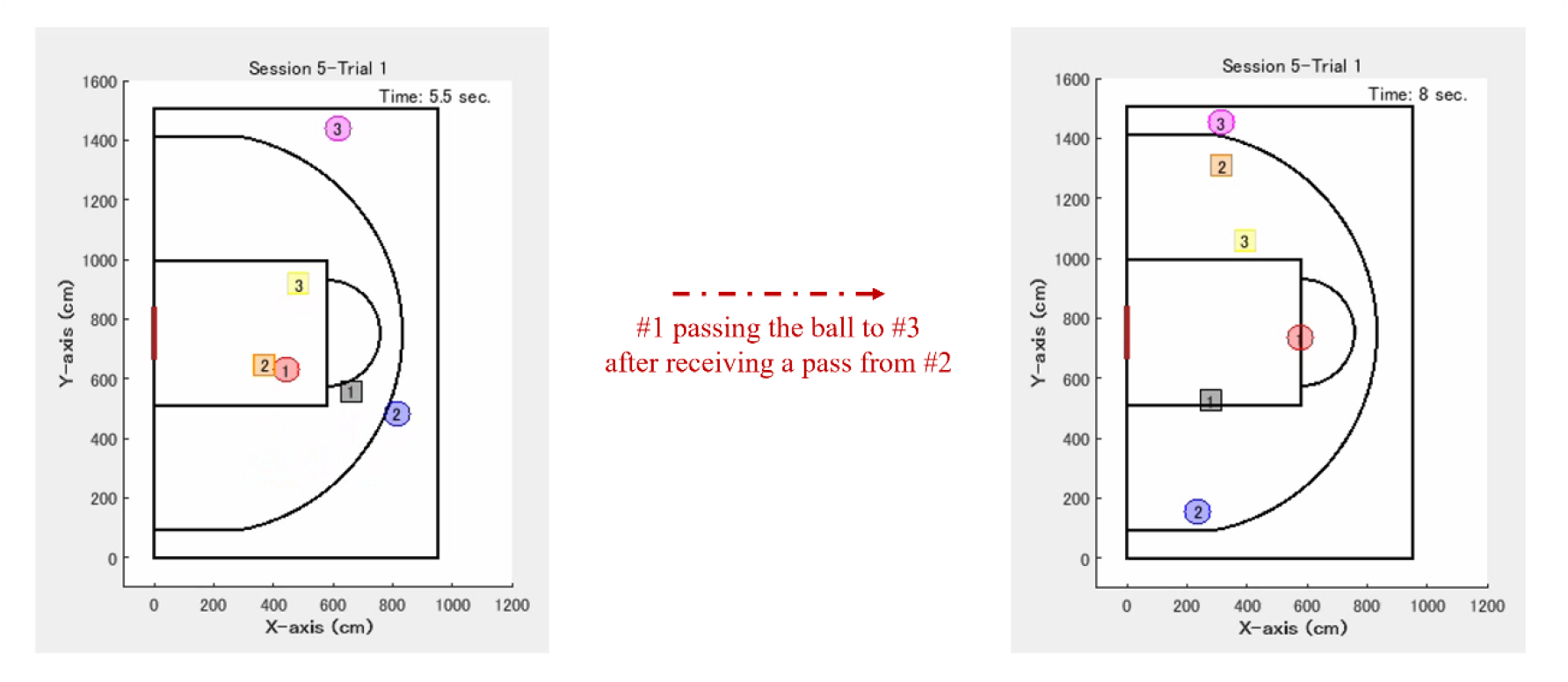
Typical example of offensive coordination after receiving the tips about the role of intervention decision and adjustment, observed in this practice. #1 runs toward the area under the basket goal (“paint area”) and receives a pass from #2. Subsequently, #3 required to this role moves to the corner area while maintaining a reasonable distance from each defensive player, receives a pass from #1, and aims to take a three-point shot.

This study may also provide the results that satisfy the usefulness and ecological validity for the role of intervention decision and adjustment. Ecological validity is related to the problem that experimental environment and observed actions significantly differ from real-world activities (51). However, after receiving the tips of this role, high offensive team performance could not be maintained in the final session (Session 6). This was caused by the defensive team establishing an effective countermeasure. After this practice, the experimenter asked the defensive team about it; the participants reported that defensive #3 marking the participant in the relevant role pretended to help the other players while actually kept marking. Thus, it was easy to establish the defensive countermeasure because the offensive pattern, as shown in Figure 8, was standardized. The offensive team had to flexibly handle the defensive one through bargaining based on the tips. However, the participants could not achieve developed coordination because of some factors, such as only a single practice and their individual skills corresponding to the third division league in Japan; they would need more training. Flexible coordination according to dynamic situations is required in various activities regardless of team sports (31, 52-54). Furthermore, both top-down and bottom-up information processing are crucial for coordination. The shared mental model of knowledge structures including strategies leads to efficient group interactions because the members are able to explain and anticipate actions each other. A group also needs to fine-tune to dynamically changing environment and others’ unanticipated actions (55-57). Defensive coordination in 5-on-5 basketball games for the top-level university team in Japan shows that the structures of role-sharing switch according to emergent situations (1). In this practice, the offensive team established shared mental model based on the tips of the crucial role. Although such top-down processing worked, developed bottom-up one might not be established in this team.

Future studies should mainly investigate (1) the number of teams participating in this practice, (2) the direct evaluation of spacing skill, and (3) these applications in other team sports. Regarding (1), only one female university team participated in this practice. The mini-games must be conducted for other teams with similar skill level. Furthermore, it is crucial to conduct the games for expert teams. The game requires basic coordinated behavior. Hence, if expert teams play it, they can express key strategies without the tips from a coach. Next as for (2), this study highlights the importance of spacing skill related to the role of intervention decision and adjustment. However, it is necessary to evaluate it directly. Similar to the previous study (50), applied geometric analysis with Voronoi diagrams enables the quantification and visualization of the occupied space of each player and the investigation of systematic offensive coordination with the player in this role. Additionally, analysis considering contexts, such as passing, dribbling, and shooting is required (e.g., (13-16)). These approaches develop deep discussions; it should be kept in mind that the results of this study indicate the overall trends of playing related to the particular role based on the experimental findings in cognitive science (21). In relation to (3), as mentioned above, spacing skill is also required in 5-on-5 basketball and soccer. Hence, if we develop a similar mini-game for other team sports, in which the relevant role is key, practical applications might be conducted. The academic and social impacts are significant because we provide an example of how to bridge the gap between controlled experiments and real-world applications. Therefore, our findings may contribute to practice design for the acquisition of spacing skill related to the relevant role and off-ball movements. Our work may offer implications for further developing 3-on-3 basketball itself. Similar to the experiment (49), a practice may be developed where the play range is limited, encouraging attention to space. For example, a coach can manipulate the offensive play range through changing the positions of defensive opponents. If a defensive player approaches to help another player, an offensive player in the role of intervention decision and adjustment role runs to a corner area and takes a three-point shot. Meanwhile, if a defensive opponent does not approach, the offensive player does not pass to this role but keeps dribbling to take a shot.

## 5 Conclusions

We applied the experimental findings in the cognitive science study of the crucial role in coordinated behavior of a triad through role-sharing to the field of sports as a pilot study. This study investigated the influence of the offensive role of intervention decision and adjustment on coordination in 3-on-3 basketball to discuss the usefulness of this role. The mini-game was developed, in which the relevant role is key, and introduced it to the practice of the female university team. The results showed that in the bins of the relatively large distance between the participant required to this role and each defensive player, the frequencies after receiving the tips of the mentioned role were significantly higher; the winning rate on the offensive team improved temporarily; however, the effects were not maintained. This suggests that spacing skill, which maintains a reasonable distance from each defensive player, emerged. Our study may provide the findings that satisfy the usefulness and ecological validity. These may bridge the gap between controlled experiments and real-world applications and contribute to practice design for the acquisition of such spacing skill and off-ball movements. In future work, it is important to examine the interactive structure to discuss the offensive coordination process.

## 7 Author Contributions

JI conceived the original concept of applying an experimental task to a 3-on-3 basketball game, designed and conducted the practice, analyzed the data, and wrote the paper. MY primarily contributed to the developing and conducting of this practice, and KF provided valuable comments throughout the research process.

## 9 Acknowledgments

We would like to thank Naruhisa Takamura, who is a coach at Beltex Shizuoka Academy, and Masanari Ichikawa of Shizuoka University for their helpful cooperation with the practice. Genki Ichinose and Yugo Takeuchi of Shizuoka University made meaningful comments; we would like to express our gratitude. This paper is published on the preprint server bioRxiv (58).

## 10 Data Availability Statement

Relevant data are provided in the Supplementary Datasets.

