## Supplementary_Material for "Analyzing coordinated group behavior through role-sharing: A pilot study in female 3-on-3 basketball with practical application"

### 1 Participants

Before this practice, we administered a profile questionnaire to the six participants. The second author, who is the head coach, divided them into the offensive and defensive teams to make their abilities competitive, referring to the results. We asked the participants about their age, dominant hand, height, basketball experience, and current and previous positions. On the offensive team, their averages were 19.33 age ($SD = 0.94$), 162.50 cm ($SD = 5.31$), and 11.33 years ($SD = 3.42$), respectively. Meanwhile, on the defensive one, they were 19.00 age ($SD = 0.82$), 167.67 cm ($SD = 5.25$), and 9.83 years ($SD = 1.31$), respectively. All the participants were right-handed. The current positions on the offensive team are the point guard, small forward, power forward, and center. Those on the defensive one are the shooting guard, small forward, and center. The previous positions on the offensive team were the point guard, shooting guard, small forward, power forward, and center. Those on the defensive one were the point guard, shooting guard, small forward, power forward, and center.

### 2 Procedures

Immediately after the first and second halves in this practice, we required each participant to answer a questionnaire. All the participants quantitatively answered questions on the degrees of (Q1) goal achievement, (Q2) collective efficacy, and (Q3) game contribution. They answered (Q1) It is easy to achieve the common goal of the winning condition on a 7-point Likert scale (0: Not well at all to 6: Very well) and (Q2) Your team can endure difficult circumstances in a game with 11-steps. -5 to +5 represent negative and positive values based on the criterion of zero before this practice. Q3 was rated from 0 to 100 under the assumption that the combined contribution to the three participants on the offensive or defensive team was 100. Japanese expression of each item was referred to in the previous studies (1,2).

The results are summarized in Supplementary Table 1. The values indicate these averages and *SD* on the offensive and defensive team. On the offensive team, the influence of the tips about the role of intervention decision and adjustment was not represented in the subjective questionnaire because only a single practice was conducted.

**Supplementary Table 1.** Results of the questionnaires in the first and second halves for the offensive and defensive teams.

| Item | Offense | | Defense | |
|---|---|---|---|---|
| | First half | Second half | First half | Second half |
| Q1: Common goal | 2.667<br>(0.943) | 3.333<br>(2.055) | 2.000<br>(1.633) | 3.667<br>(0.943) |
| Q2: Collective efficacy | 1.000<br>(0.816) | 1.667<br>(2.357) | 0.333<br>(1.700) | 2.667<br>(0.471) |
| Q3: Game contribution | 53.333<br>(20.548) | 51.667<br>(13.123) | 43.333<br>(28.674) | 43.333<br>(20.548) |

a Value was the average in each team. Parentheses indicate these *SD*s.


### 3 Results

This study analyzed (1) the distance (cm) between the offensive participant required to the role of intervention decision and adjustment, as shown in pink in Figure 1, and each defensive player, as shown in black, orange, or yellow in Figure 1, and (2) that between the offensive key player and each other participant, as shown in red or blue in Figure 1. We calculated both indices of (1) and (2) for each time frame and made these histograms. The histograms were made in each trial and the normalized frequencies were averaged for the first and second halves. A $t$-test was conducted to compare them between the first and second halves in each bin and an effect size (Cohen's $d$) and power ($1-\beta$) were calculated. Supplementary Figure 1 and Figure 2 show all the averages of the normalized frequencies in each bin for the indices of (1) and (2). Supplementary Datasets indicate these analysis results; you can see the details on the *t*-value, degree of freedom, *p*-value, effect size, and power indices, including cases where no significant differences were found.

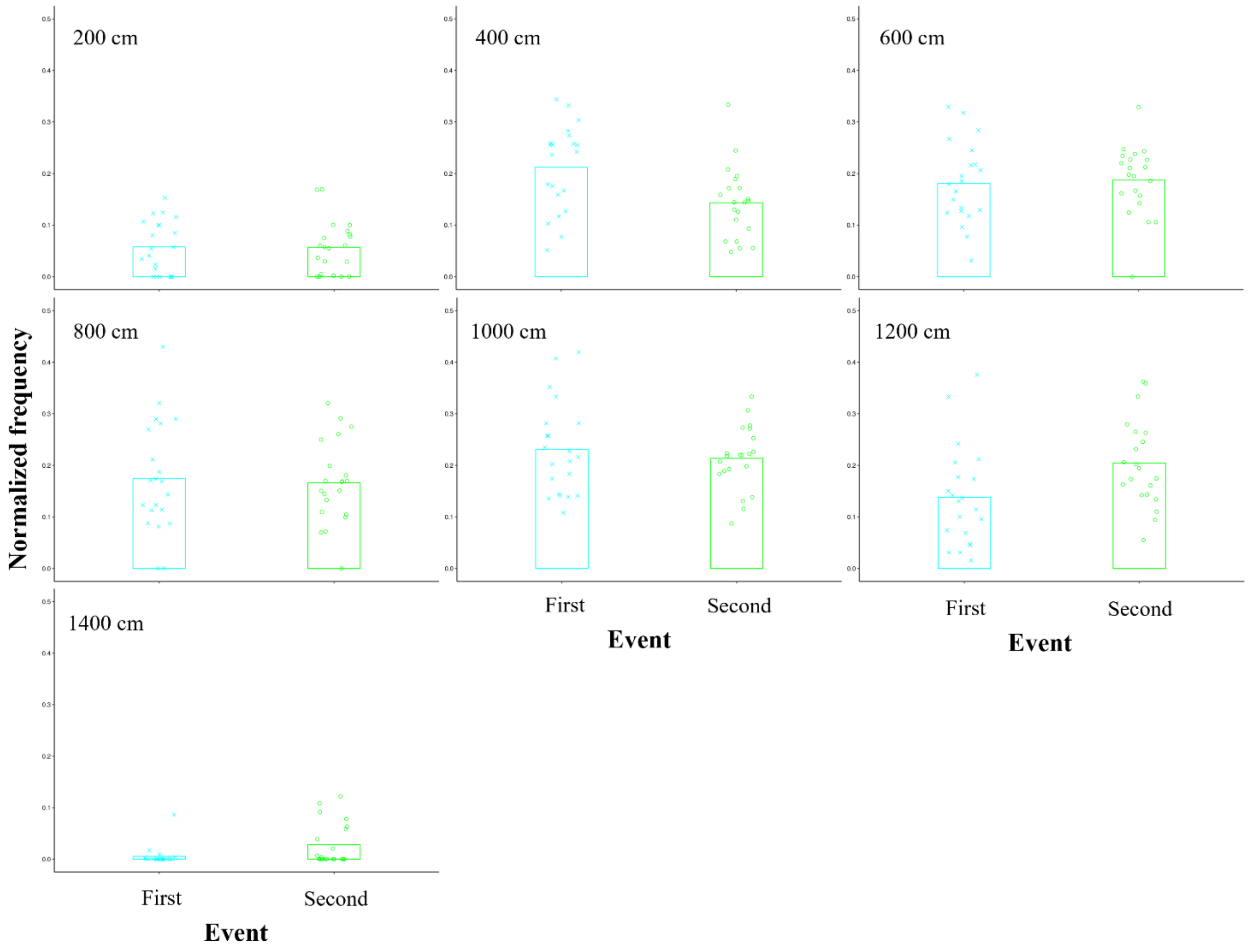


**Supplementary Figure 1.** Averages of the normalized frequencies in all the bins for the index of the distance (cm) between the offensive player required to the role of intervention decision and adjustment and each defensive participant. The range of the bins is from 200 cm to 1400 cm.

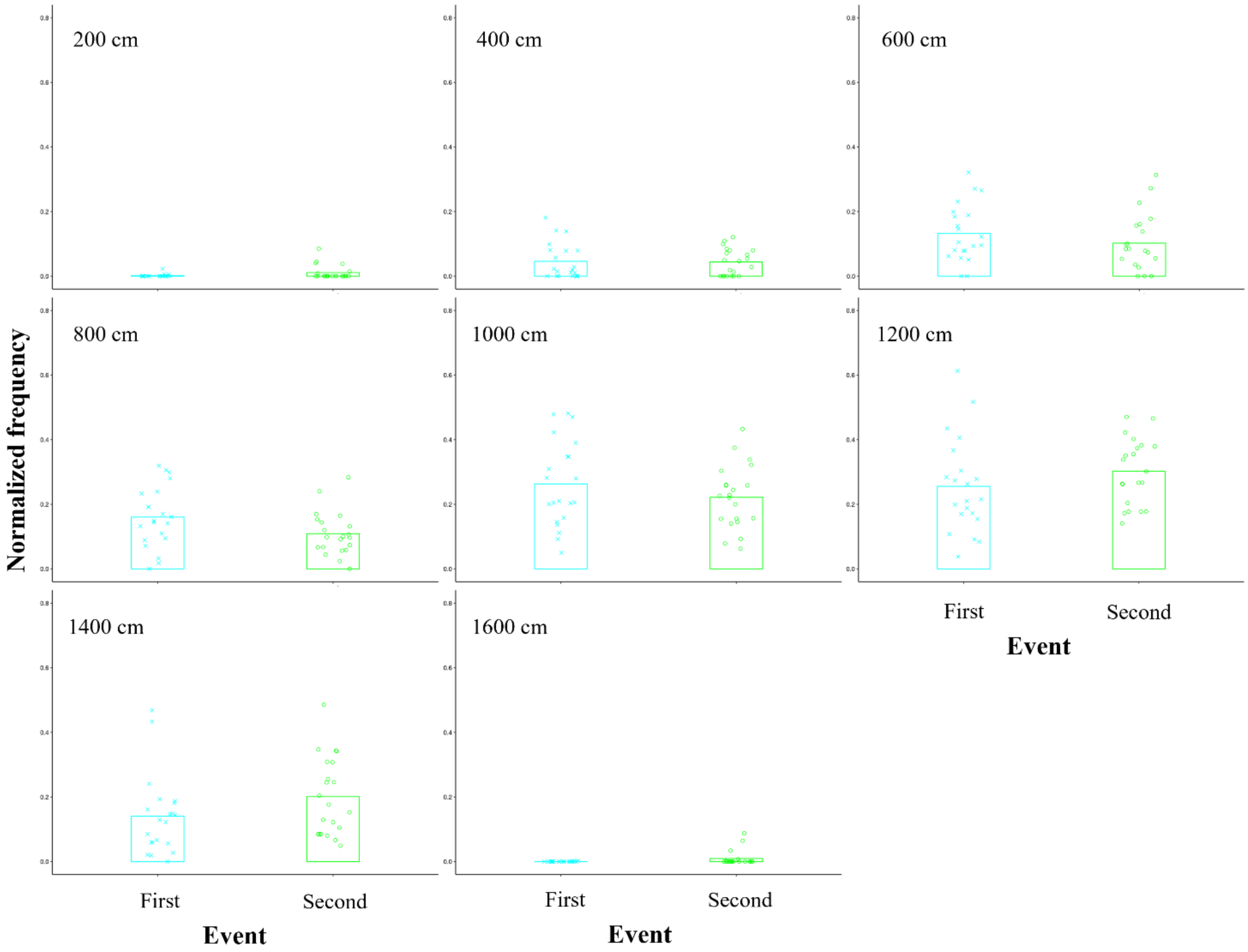


**Supplementary Figure 2.** Averages of the normalized frequencies in all the bins for the index of the distance (cm) between the offensive player required to the role of intervention decision and adjustment and each other participant. The range of the bins is from 200 cm to 1600 cm.
